# Biofilm lifestyle as a common trait of ammonia-oxidizing archaea

**DOI:** 10.1101/2024.11.18.624116

**Authors:** Maximilian Dreer, Thomas Pribasnig, Logan H. Hodgskiss, Zhen-Hao Luo, Fran Pozaric, Christa Schleper

## Abstract

Although widespread in nature, growth in biofilms has been relatively little explored in the globally distributed ammonia oxidizing archaea (AOA). Here we investigated six representatives of three different terrestrial and marine clades of AOA in a longitudinal and quantitative study for their ability to form biofilm and studied gene expression patterns of three representatives. While all strains grew on a solid surface, soil strains exhibited the highest capacity for biofilm formation. Based on microscopic and gene expression data, two different colonization strategies could be distinguished. S-layer containing AOA (from both soil and marine habitats) initialized attachment as single cells and subsequently formed denser layers and three-dimensional structures, while the S-layer free species of the *Nitrosocosmicus* clade attached as suspended aggregates to the surface and henceforth showed fastest establishment of biofilm. Transcription profiles were significantly different between planktonic and biofilm growth in all strains and revealed individual reactions, often fulfilling shared functions. In particular the strong expression of different types of multicopper oxidases was observed in all strains indicating modifications of their cell coats. S-layer carrying AOA each additionally expressed a set of adhesion proteins supporting attachment. Detoxification of nitrous compounds, copper acquisition as well as the expression of transcription factor B were also shared reactions among biofilm producing strains. However, the majority of differentially expressed protein families was distinct among the three strains illustrating that individual solutions have evolved for the shared growth mode of biofilm formation in AOA, probably driven by the different ecological niches.

## Introduction

Nitrification, the microbially mediated oxidation of ammonia (NH_3_) to nitrate (NO_3_^-^) via nitrite (NO_2_^-^), is a key process of the global biogeochemical nitrogen cycle. The first and rate-limiting step of ammonia oxidation (NH_3_ to NO_2_^-^) is performed by ammonia oxidizing archaea (AOA), ammonia oxidizing bacteria (AOB) and comammox, capable of complete oxidation of NH_3_ to NO_3_ ^-^. All ammonia oxidizing microorganisms contribute directly or indirectly to the production of nitrous oxide^1,2^, a potent greenhouse gas, and to the loss of nitrogen in natural and in fertilized agricultural systems^3,4^. It is therefore of continuous importance to better understand their metabolisms and activities in diverse ecosystems. AOA are ubiquitous in most aerobic environments, ranging from the sediments of the Mariana trench ^5^ to the soils of Mount Everest^6^. They outnumber their bacterial counterparts in most oligotrophic environments including pristine soils and the open ocean^7,8^ often orders of magnitude. Although physiological studies have indicated a specialization to low nutrient environments^9,10^, AOA populations additionally outnumber AOB in highly fertilized agricultural soils^7^, indicating so far unknown metabolic capabilities or a wider range of growth modes than currently known.

AOA first belonged to the class Nitrososphaeria^11^ in the phylum Thaumarchaeota^12^ but have since been reclassified into the order Nitrososphaerales including the three major families “*Candidatus* Nitrosocaldaceae”, *Nitrosopumilaceae*, and *Nitrososphaeraceae*^13^. While most of the isolated AOA strains from aquatic or terrestrial environments have been isolated in liquid media^14,15^, some species from soil, in particular those of the *Nitrosocosmicus* clade were reported to grow on solid surfaces^16–18^, or as aggregates in liquid culture^19^, indicating the capacity to grow in biofilms. *Ca*. Nitrosocosmicus oleophilus MY3 even showed increased growth rates when attaching to surfaces^16^. Interestingly, *Ca*. Nitrosocosmicus are also present in low abundances in wastewater treatment plant biofilms dominated by thecomammox *Nitrospira*, dominate these communities^20,21^. Despite those observations, a systematic insight into the capability of biofilm formation of different AOA is lacking as is a fundamental understanding of their physiology under these conditions.

It has been hypothesized that most microorganisms form or are a part of biofilms in-situ. With estimates of 40-80% of cells on earth residing in biofilms, they are also hypothesized to drive all biogeochemical cycles^22^. While most soil microorganisms are thought to grow in biofilms, marine organisms can grow planktonically or particle associated, offering specialized niches for different species^23^. Biofilms are defined by the IUPAC as “aggregates of microorganisms in which cells are frequently embedded in a self-produced matrix of extracellular polymeric substances (EPS) that are adherent to each other and/or a surface”^24^. This is in line with previous and broad definitions of biofilms, including microbial aggregates, floccules and also adherent populations on surfaces, in contrast to single planktonic or single sessile cells^25^. Additionally, biofilms can have so-called emergent properties, differing substantially from the physiology of free-living cells. Those properties arise mostly from the EPS-matrix the cells are thriving in, and include but are not limited to: protection against biotic and abiotic stress, sorption of nutrients, retention of enzymes and localization of gradients^26^. The EPS-matrix is typically mainly made of extracellular polysaccharides, proteins, lipids and DNA, however compositions vary between species^27^.

We hypothesize that biofilm formation is a general trait of AOA and that general physiological changes can be identified in this potentially ecologically relevant mode of growth. The propensity of six representatives, covering major lineages of AOA, to form biofilm was studied and determined by a growth assay using borosilicate glass as a surface. Strains included were AOA isolated from garden, arable and acidic arable soil (*Nitrososphaera viennensis* EN76, *Ca.* Nitrosocosmicus franklandianus C13 and Nitrosotalea sinensis Nd2, respectively), tropical marine aquarium gravel (*Nitrosopumilus maritimus* SCM1) and coastal surface water (*Nitrosopumilus piranensis* D3C*, Nitrosopumilus adriaticus* NF5)^15,19,28,14,29^. A genome wide transcriptomic analysis comparing planktonic growth to growth in a biofilm of three representatives, *N. viennensis*, *Ca*. N. franklandianus (hereafter referred to as *N. franklandianus*) and *N. maritimus* was performed and is presented along with SEM and light microscopy imaging, including a time series elucidating the architecture and succession of AOA biofilms in individual strains using microscopic techniques.

## Material and Methods

Detailed cultivation description, culture maintenance, protocols for RNA extraction, transcriptomic analysis and phylogenetic tree calculation are available in the supplementary material.

### Growth as biofilm

Borosilicate cover glasses (CG) (VWR^TM^, 631-0120, 18x18 mm) or soda-lime glass microscope slides (MS) with a silanized surface leading to a positive charge (Carl Roth®, Histobond®, CEX0.1, 76x26x1 mm) were sterilized by autoclaving in Milli-Q® water, after which they were individually added to 30 ml polystyrene containers or 250 ml schott bottles containing 20 ml or 125 ml growth media respectively using tweezers sterilized with Incidin™. The growth media were inoculated and grown as described in supplementary material and are summarized in Table S1. Upon reaching a predefined range of nitrite (Table S1), the CG or MS were transferred to fresh growth media using sterilized tweezers. To ensure that only cells attached to the provided surfaces were transferred, excess liquid forming droplets at the edges of CG was absorbed using UV sterilized whatman paper (Whatman™ GB003). MS were carefully dipped three times in prewarmed basal media before being transferred to fresh medium. Using this setup CG and MS were continuously transferred. Nitrite was measured and cultures checked for purity as described in supplementary materials for stock cultures. Biofilm cultures were never inverted before sampling to prevent the disruption of the biofilm. CG were fully submerged standing upright and slightly tilted at the bottom of the 30 ml containers due to the container’s internal diameter of 22.38 mm. MS were not fully submerged, the frosted area not being covered, standing upright and strongly tilted in 250 ml schott bottles. After reaching the species-specific fastest nitrite production, five MS each of *N. viennensis*, *N. franklandianus* and *N. maritimus* biofilms were frozen on dry ice and stored at -70°C for RNA extraction.

### Microscopy of biofilms

To directly image undisturbed in-situ biofilm on CG, imaging spacers from an adhesive sheet with a thickness of 0.12 mm (Grace Bio-Labs SecureSeal^TM^ adhesive sheets) were prepared. The sheet was cut into CG sized frames with an inner dimension of ∼13x13 mm, attached to microscopy slides and 40µl of basal medium (FWM or SCM) added to the middle of the frame. Active CG biofilms were immediately mounted to the prepared frames using tweezers after taking off excess liquid as described above. The CG were sealed with nail polish, dried for 15 minutes in the dark at room temperature and imaged with phase contrast microscopy.

For scanning electron microscopy CG were prepared as follows. After initial in-situ fixation of CG in growth media with 2.5% glutaraldehyde for 5 min at room temperature, CG were transferred to PBS buffer containing 2.5% glutaraldehyde and stored at 4°C overnight. The CG were washed three times in PBS before being dehydrated in an ethanol series (30%, 50%, 70%, 80%, 90%, 100%) and dried via either critical point drying with CO_2_ (*N. viennensis*, *N. maritimus*), or chemically using Hexamethyldisilazane (*N. franklandianus*). Dried samples were sputter coated with Au and imaged in a JEOL IT 300 scanning electron microscope at 20 kV. To prevent the disruption of biofilm structures CG were standing upright throughout the whole process using the 30 ml polystyrene containers for initial fixation, 50 ml falcon tubes for fixation at 4°C and dehydration and the cover glass holder of the Leica EM CPD300 for critical point drying.

## Results

### Biofilm formation in different AOA

AOA were initially grown planktonically in the presence of a borosilicate cover glass (CG) as a surface for attachment before consecutive transfers of only the CG and attached cells. The ability of AOA to attach to CG and accumulate biomass over time was investigated by following nitrite production, which is a well established proxy for cell numbers^14,15^ (Fig. 1, Dataset S1). In this experimental setup, nitrite production did not exclusively stem from biofilms, as a planktonic fraction was observed for every transfer. However, only cells adhering to cover glass were transferred, implying that the planktonic fraction in the following cultures was seeded from biofilms. A faster rate of nitrite production should therefore represent an increased amount of biomass on the cover glass. The rate of nitrite production of all tested AOA increased over the course of multiple CG transfers indicating the capacity for biofilm formation.

**Figure 1.**
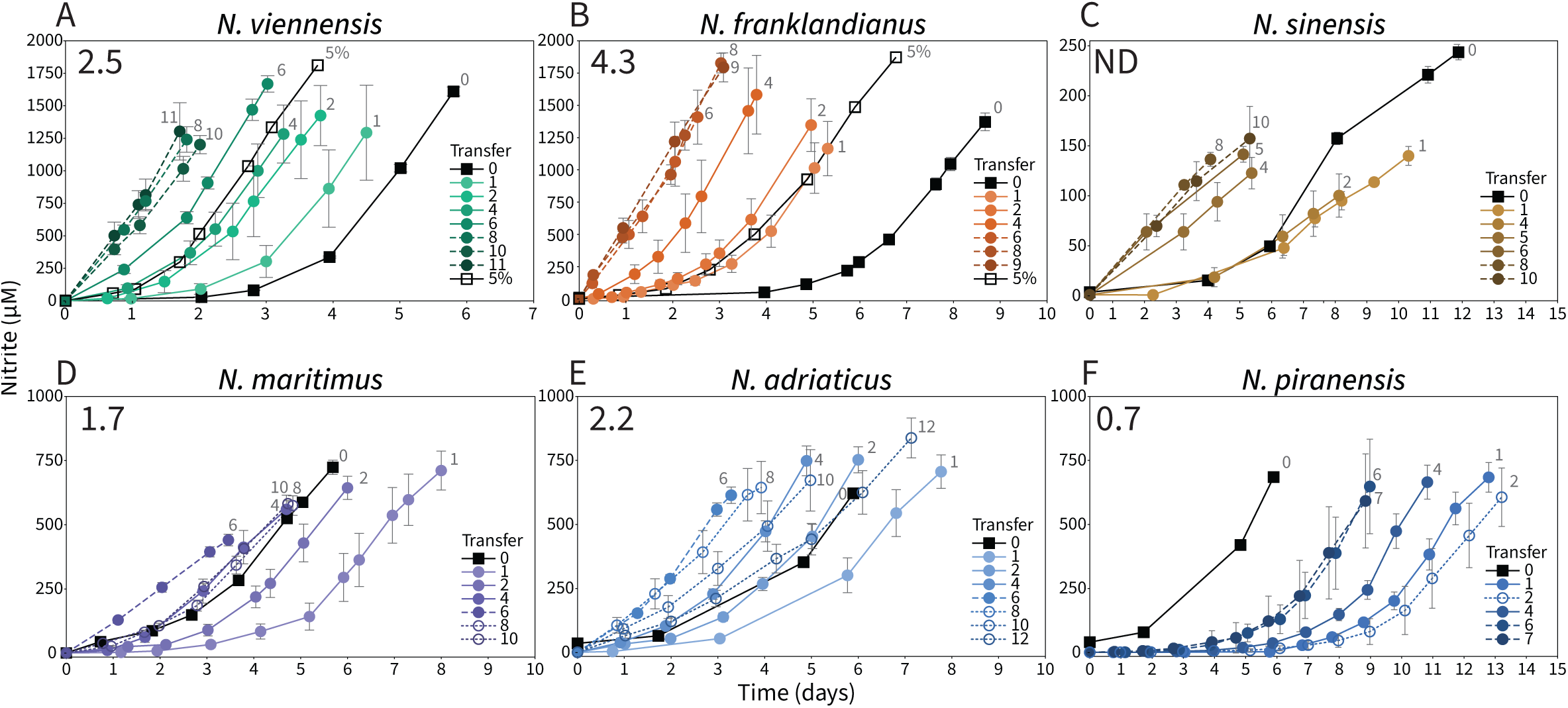
Biofilm formation of ammonia oxidizing archaea on cover glass (CG). Nitrite production of (A) *Nitrososphaera viennensis,* (B) *Nitrosocosmicus franklandianus*, (C) Ca. *Nitrosotenuis sinensis*, (D) *Nitrosopumilus maritimus*, (E) *Nitrosopumilus adriaticus* and (F) *Nitrosopumilus piranensis* grown as biofilm on cover glass. Initial planktonic growth of cells in the presence of CG (black lines), continuous transfers of CG (colored lines), and maximum nitrite production rate (colored dashed lines) are shown. Drops in nitrite production rates between transfers were observed for marine strains (dotted lines). Cultures started with 5% inoculation volume done in duplicate are shown for *N. viennensis* and *N. franklandianus* (black line, empty rectangle). Maximum BARs values are displayed for each species. Increasing number of transfers are indicated by numbers next to lines and a darkening color gradient. Nitrite measurements show averages of eight biological replicates. Error bars depict the standard deviation. For illustrative purposes not all transfers are depicted. The raw data of all transfers can be found in Dataset S1.

The soil AOA *N. viennensis* and *N. franklandianus* exhibited a clear increase in nitrite production rate already after the initial cover glass transfer that continuously accelerated with each consecutive transfer (Fig. 1A, B). A stable maximum for both species was reached after 8 and 6 transfers respectively (Fig. 1A, B). Similarly, *N. sinensis* reached a stable maximum after 8 transfers well above the nitrite production of initial growth, however, its growth was considerably slower than the two soil strains and not exponential. The nitrite production rate of all marine AOA strains was less stable with successive CG transfers indicating a less pronounced capability of biofilm formation (Fig. 1D-F). After peaking at the sixth transfer, the nitrite production rate of *N. maritimus* decreased and stabilized at a lower rate, while it steadily declined in *N. adriaticus*. Unlike the other marine strains, the nitrite production rate of *N. piranensis* started to decrease already with the first two transfers before eventually showing an increase in nitrite production rate. This along with its high standard deviations in successive CG transfers suggests the lowest capability for biofilm formation as compared to all other strains (Fig. 1F).

The stark differences between the initial growth and first transfer of the *Nitrososphaeraceae* and *Nitrosopumilaceae* species can be attributed to the different inoculation volumes used, ranging from 0.25% to 5% (v/v). These inoculation volumes were chosen in order to transfer the CG of all species after an equal period of time given for the initial transfer. A line displaying growth of a 5% (v/v) inoculation culture was added for *N. viennensis* and *N. franklandianus* as reference.

To be able to quantitatively compare the biofilm forming capabilities of different strains, a biomass accumulation rate (BAR) was calculated - taking species-specific generation times into account. The time required to produce 500 µM of nitrite was determined for both, planktonic 5% inoculation volume cultures and all CG transfers and termed standard time (ST) and biofilm time (BT) respectively. The BAR was then calculated by dividing the ST by the BT for each transfer, expressed as BAR = ST/BT_x_, where *x* represents the transfer number.

The maximum BARs support that *N. franklandianus* had the highest biofilm-forming capabilities. Although the marine strains *N. maritimus* and *N. adriaticus* initially displayed biofilm-forming abilities comparable to those of *N. viennensis*, they were not stable over time (Fig. 1 and Table S2). Since *N. sinensis* did not exhibit exponential growth, it was excluded from the analysis.

### Time series of biofilm formation

The in-situ biofilm formation on CG by *N. viennensis*, *N. franklandianus* and *N. maritimus* was imaged for five consecutive transfers by light microscopy. After initial growth in the presence of CG, single or small aggregates of attached cells were observed (Fig. 2-T1). Over the course of the two following transfers microcolonies and larger tower-like structures formed and continued to expand (Fig. 2-T2 - T4). These structures eventually merged, establishing either multilayered, three-dimensional biofilms in *N. viennensis* and *N. franklandianus*, or a monolayer with small, interspersed aggregations in *N. maritimus* (Fig. 2- T3 - T4). The biofilm of all species eventually covered the whole surface of the CG in a continuous or interspersed layer (Fig. 2-T5).

**Figure 2.**
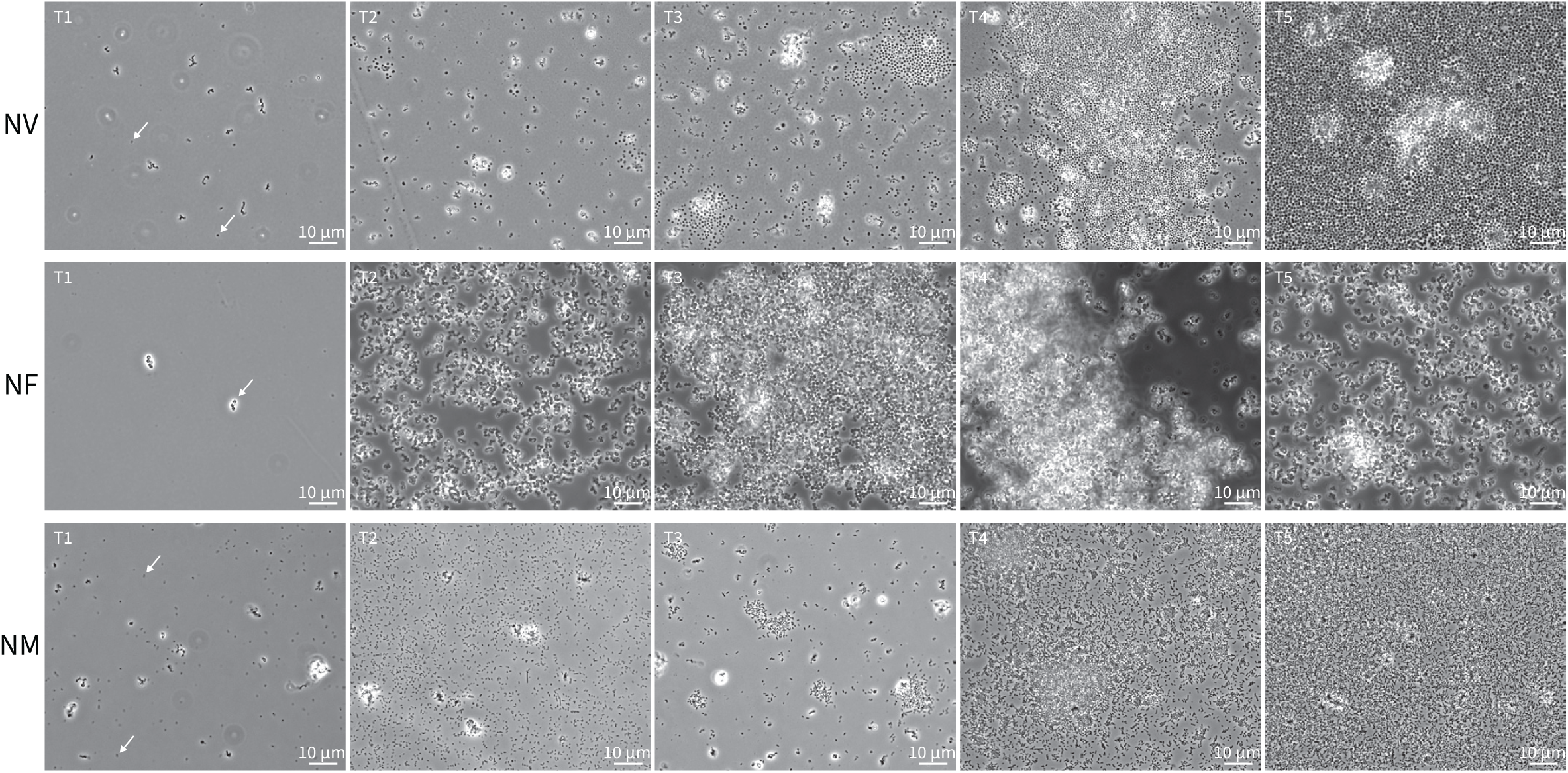
Light microscopy time series of *N. viennensis* (NV) *N. franklandianus* (NF) and *N. maritimus* (NM) in-situ biofilm formation on cover glass (CG). For five transfers CG were destructively sampled for light microscopy imaging after production of 1200- 1600µM NO_2_. (T1) Initial growth in presence of a CG. (T2-5) First to fifth transfer of CG. White arrows indicate cells initially adhering to a CG (T1). Observed features included: multicellular structures/microcolonies (T2, T3). Three-dimensional, multilayered structures (NF-T3, T4, T5, features out of focus appeared white and blurry). Biofilm fully covering the CG (NF-T3/T4, T5).

Two differing colonization strategies were identified: *N. viennensis* and *N. maritimus* attached to the CG as single cells, which formed microcolonies that subsequently merged into more extensive three-dimensional structures. The attachment of *N. franklandianus* on the other hand appeared to be based on the deposition of small, pre-formed suspended aggregates and happened much more rapidly, with three-dimensional tower-like structures already forming from the second transfer on. Along with its rapid biofilm development, *N. franklandianus* also formed the most extensive, multilayered biofilms, characterized by the height and area of the observed tower-like structures. (Fig. 2-NF-T3,4).

### Different morphologies of AOA biofilms

For scanning electron microscopy (SEM) of *N. viennensis*, *N. franklandianus* and *N. maritimus* biofilms, the CG shown in Fig. 1 were prepared without removing the biofilm once stable nitrite production maxima were reached. SEM not only confirmed the multilayered and three-dimensional nature of biofilms formed by *N. viennensis* and *N. franklandianus*, but also revealed the presence of putative EPS in the biofilms formed by both organisms (Fig. 3A, B, D, E) In *N. viennensis*, EPS formed thread-like structures that linked individual cells or clusters of cells together, while *N. franklandianus* exhibited granular EPS clusters between closely associated cells. *N. maritimus* biofilm structures were expectedly less extensive, but displayed a surprising amount of putative EPS (Fig. 3C, F) consisting of both an extensive granular EPS scaffold and minimal thread-like structures connecting few cells.

**Figure 3.**
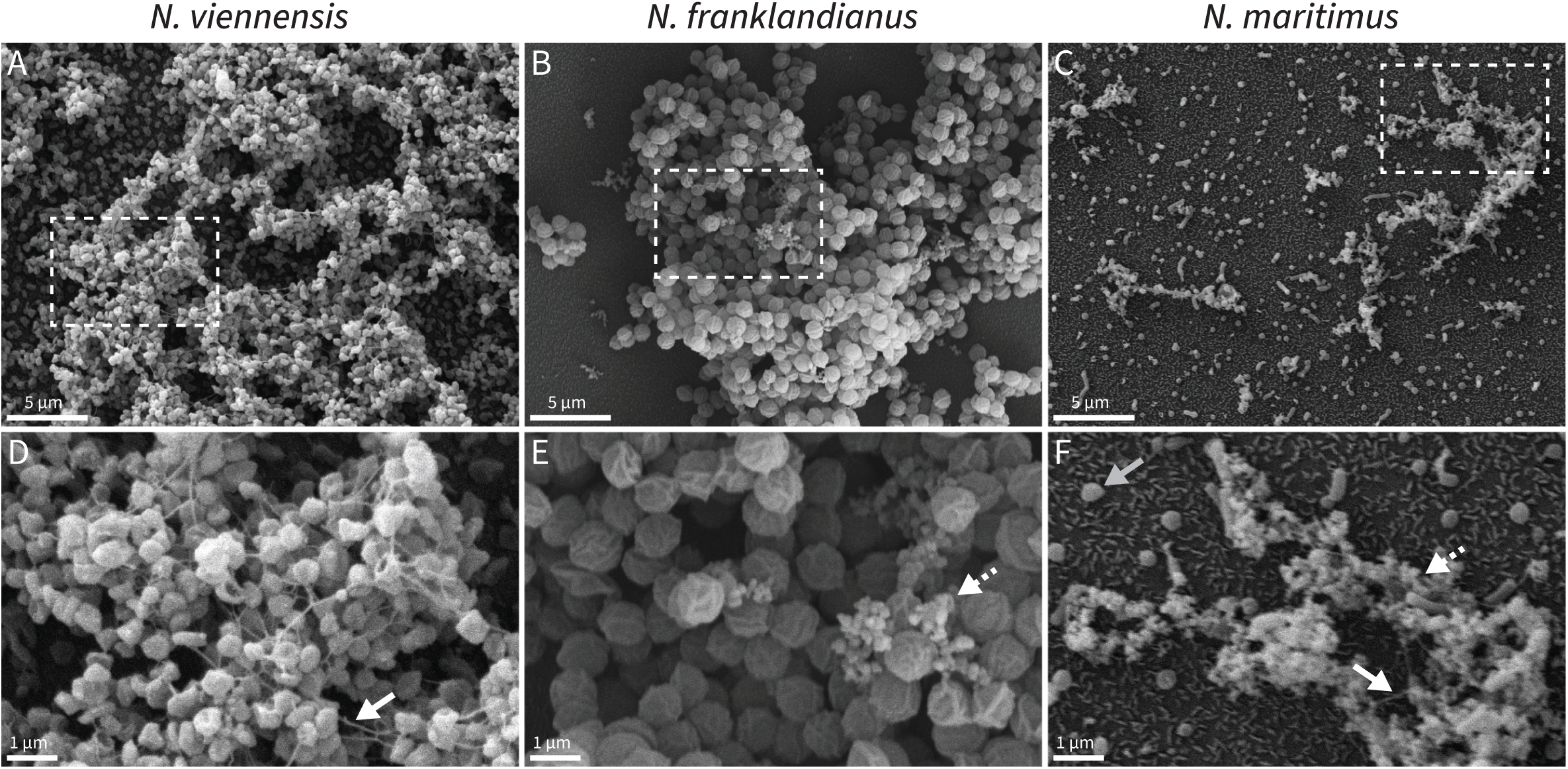
Visualization of AOA biofilms by scanning electron microscopy (SEM). *N. viennensis* (A, D) *N. franklandianus* (B, E) and *N. maritimus* (D, F). Putative EPS were detected as thread-like structures for *N. viennensis* and *N. maritimus* (D, white arrows) and granular structures for *N. franklandianus* and *N. maritimus* (E, F, white dashed arrows). *N. maritimus* cells collapsed due to SEM preparation were visible (F, gray arrow). Micrographs D-F are enlargements of areas marked with white dashed boxes in A -C respectively.

### Biofilms reveal a distinct transcriptional state in each species

To further characterize the differences of growth in biofilm to planktonic conditions, transcriptomic analyses were done for *N. viennensis*, *N. franklandianus* and *N. maritimus* (Datasets S2-4). Genes with a log_2_FC of >1.0/< -1.0 and an adjusted p-value cut-off of 0.001 were considered significantly up- or downregulated. In *N*. *viennensis, N. franklandianus* and *N. maritimus* 250/3187, 135/2768, 104/1972 genes were significantly upregulated and 59/3187, 20/2768 and 51/1972 genes were significantly downregulated, respectively. PCA analyses revealed distinct transcriptomic states, clearly separating the different conditions on PC1 (Fig. 4). Variation between biological replicates of the same condition displayed on PC2 was more pronounced in biofilms than the planktonic controls. When comparing the 25 strongest upregulated gene families (Fig. 4) only few responses were found to be conserved among the transcriptomes of the three strains. Those included multicopper oxidases (MCOs) and the archaeal transcription initiation factor (TFB). Most other highly upregulated genes within each strain were predominantly species-specific, with a slightly larger overlap among the soil strains (Fig. 4, “Upreg. BF”).

**Figure 4.**
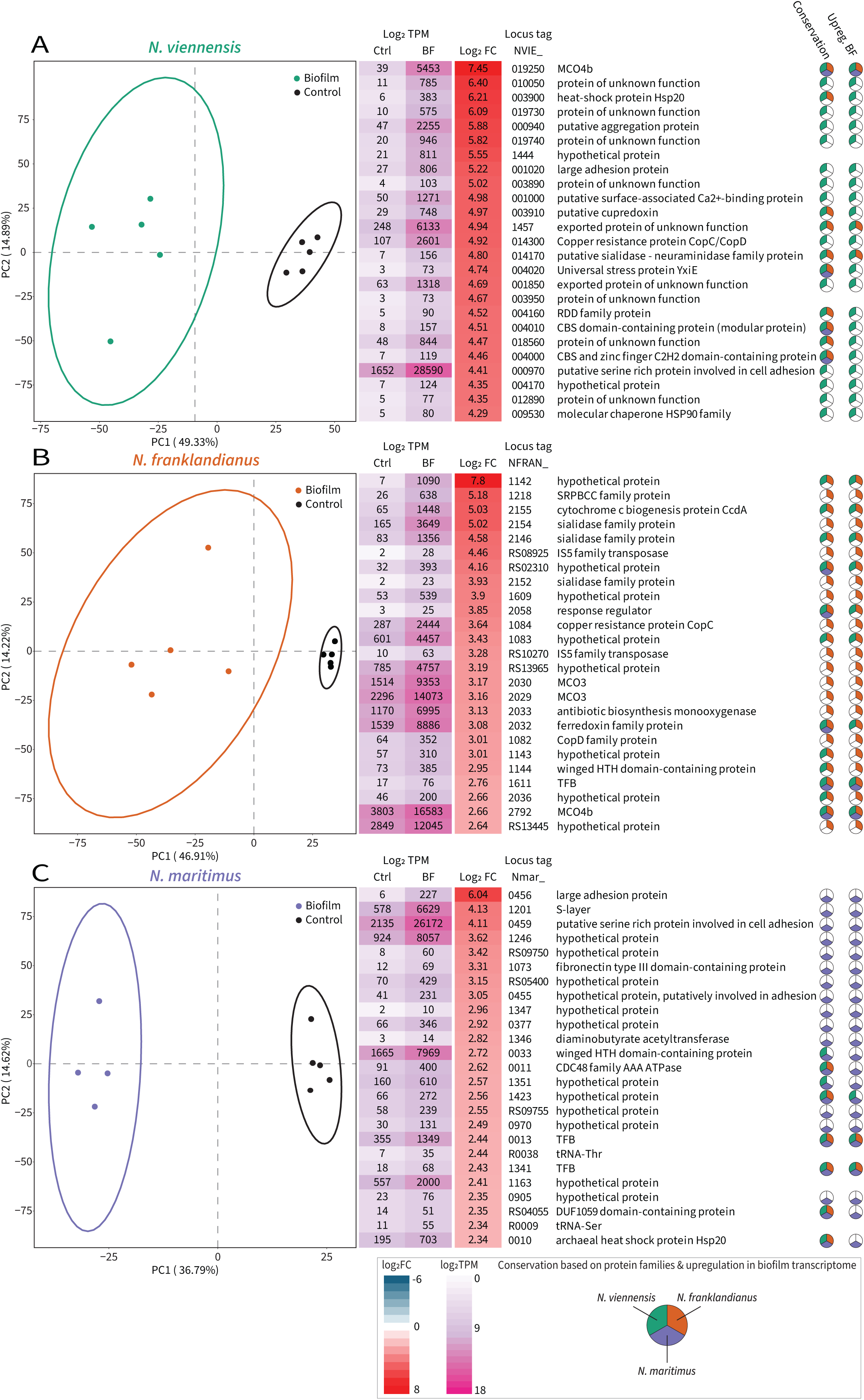
PCA plots & Top25 upregulated genes: Principal component analysis of log transformed expression data. Principal component 1 (PC1), explaining between 36.79%- 49.33% of variance clearly separates conditions from each other, while principal component 2 (PC2), including 14.22%-14.89% of variance, describes the variance within conditions. Ellipses represent a confidence interval of 95%. PERMANOVA analysis indicated a significant difference between BF and Ctrl conditions based on the first two principal components (NV: p = 0.002; NF: p =0.008; NM: p =0.009, cutoff =0.01). Top 25 differentially expressed genes are ordered by log_2_FoldChange. TPM, expressing levels of transcripts per million, within the transcriptome are displayed for both planktonic controls (Ctrl) and biofilms (BF) and were color coded after log_2_ transformation (Datasets S2- S4). Wedged circles on the right indicate the genes genomic conservation and upregulation in biofilms in NV (green), NF (brown) or NM (purple)

### Few Conserved Gene Families Define Biofilm Formation Across Species

The analysis was expanded to identify overlaps in upregulated protein families of all genes considered significantly upregulated. The majority of genes were linked to a protein family upregulated in only one out of each of the three species (195 in *N. viennensis*, 102 in *N. franklandianus* and 83 in *N. maritimus*) (Fig. S1), emphasizing their species-specific responses. Out of all protein families, only three were significantly upregulated in biofilms of all three species (Fig. S1, Dataset S5). Of these, proteins of the multicopper oxidase family (MCO) were among the most differentially expressed in *N. viennensis* and *N. franklandianus*. A phylogenetic tree expanding the earlier work of Kerou et al. to include 143 AOA species was reconstructed and used to differentiate the upregulated MCOs potentially involved in biofilm formation (Fig. S2)^30^. A simplified version including only MCOs of the species included in this study is shown below (Fig. 5).

**Figure 5:**
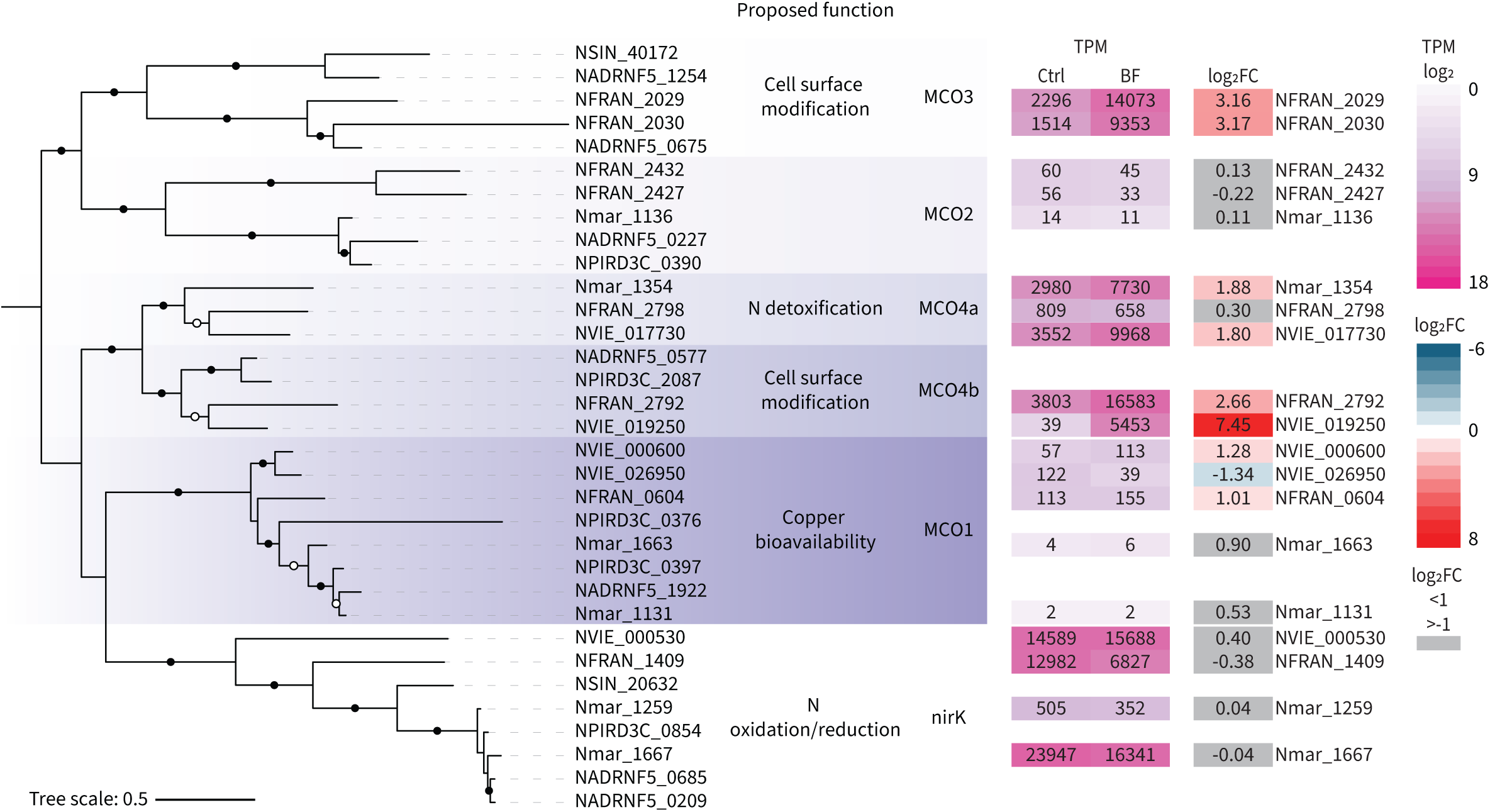
Phylogenetic tree of MCOs in *N. viennensis*, *N. franklandianus, N. sinensis*, *N. maritimus, N. adriaticus* and *N. piranensis*. Significant and not significant log_2_-FC are displayed and colored as gradient or gray respectively. TPM displayed for planktonic controls (Ctrl) and biofilms (BF) were color coded after log_2_ transformed TPM (Datasets S2- 4).

While MCO1 and MCO2 genes were not highly transcribed in either planktonic conditions or biofilms, several MCO4 and MCO3 genes were found to be upregulated in biofilms. MCO4 genes can be further divided into two subtypes. The first subtype, MCO4a, was highly transcribed in planktonic conditions and further upregulated in biofilms of *N. viennensis* (NVIE_017730) and *N. maritimus* (Nmar_1354). In *N. franklandianus*, however, this subtype (NFRAN_2798) was only moderately transcribed in both conditions. The two soil strains harbor an additional subtype MCO4b, which is not present in *N. maritimus*. MCO4b was the most upregulated gene in *N. viennensis* (NVIE_019250) and also highly upregulated in *N. franklandianus* (NFRAN_2792). Additionally, *N. franklandianus* contains two copies (NFRAN_2029, NFRAN_2030) of type MCO3, which is not present in either *N. viennensis* or *N. maritimus*. Both MCO3s were among the most upregulated genes in biofilms of *N. franklandianus*.

The second protein family found upregulated in all three species was the archaeal transcription factor B (TFB), guiding the initiation of transcription in archaea^31^. At least one TFB gene was upregulated in all three species. In *N. viennensis*, the highest expressed TFB (NVIE_012290) was further upregulated, while the expression of the other four TFBs remained unchanged (Dataset S6). In contrast, in *N. franklandianus*, no single TFB dominated, but three different TFBs (NFRAN_2995, NFRAN_3010, NFRAN_3143) were expressed equally under both conditions, while an upregulation of three other, less expressed TFBs (NFRAN_0924, NFRAN_1611, NFRAN_2944) was observed. Only *N. maritimus* (which contains eight TFBs) clearly shifted its highest expressed TFB from Nmar_0517 in planktonic conditions to Nmar_0013 in biofilms (Dataset S6).

Finally, a putative nitroreductase was upregulated in all three species, although the differential expression was not as prominent with log_2_ fold changes between 1-1.35 and low expression levels (Dataset S6).

Unsurprisingly, the overlap of upregulated genes between the two soil strains was bigger than any other overlap, and included sialidases, genes involved in urea transport or utilization and several regulatory genes (Datasets S5, S6).

### Expression patterns and gene clusters of *bona fide* biofilm genes

Under the assumption that genes important under biofilm conditions should be upregulated and highly expressed in biofilms, a specific set of biofilm-associated genes was extracted (Fig. 6). Firstly, the top 50 upregulated genes of every species were filtered by top 100 TPM to identify genes highly upregulated <u>and</u> highly expressed in biofilms. Secondly, functionally related genes in close proximity to the identified upregulated genes and following the same expression pattern, but not the chosen cutoffs, were also included (marked by an asterisk in Fig. 6).

**Figure 6.**
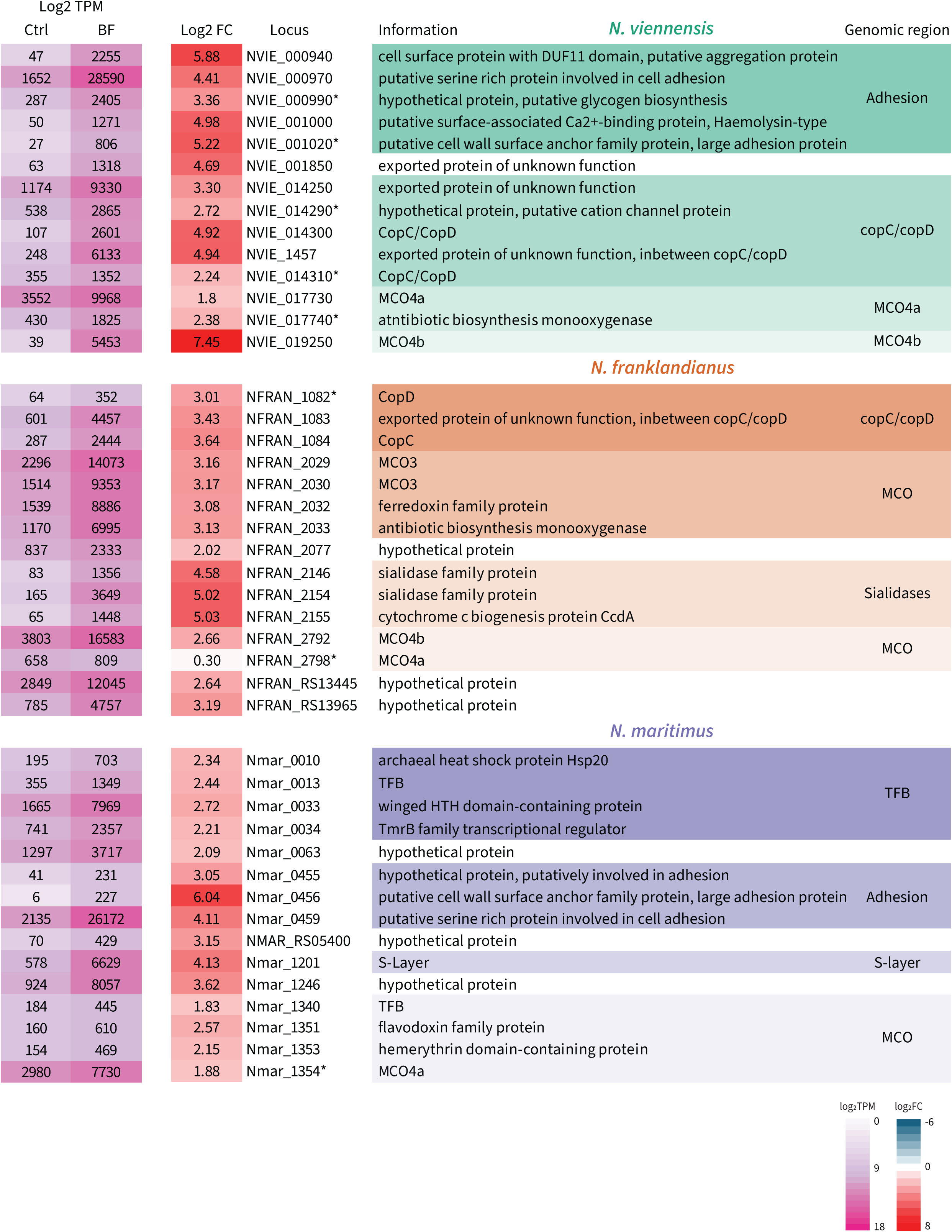
Bona fide biofilm genes. Top 50 log_2_FC biofilm genes of *N. viennensis*, *N. franklandianus and N. maritimus* were filtered by TPM rank < 100 to define genes that are highly upregulated and highly expressed in AOA biofilms. The locus tag of manually added genes were marked with asterisks (*). These genes were either not included in the top 50 log_2_FC or TPM rank < 100 biofilm genes, but were found in the direct vicinity of bona fide biofilm genes and followed the same general expression pattern. Shaded regions indicate genes with putative functional similarity and relative synteny (20 genes or less apart). Genes with “RS” indicate genes with a RefSeq annotation but no corresponding GenBank annotation.

The following patterns emerged:

Several genomic regions of biofilm associated genes were identified, especially in the soil strains. In contrast, less regions were identified and the biofilm-associated genes were more dispersed across the genome of *N. maritimus*. (Fig. 6)

In *N. viennensis* and *N. franklandianus,* several MCOs were highly upregulated and highly expressed. Both MCO4a in *N. viennensis* and MCO3 in *N. franklandianus* clustered with a gene annotated as antibiotic biosynthesis monooxygenase. In *N. franklandianus* a ferredoxin family protein gene was also part of this region. In N. *maritimus*, a hemerythrin and a flavodoxin family protein gene clustered with MCO4a and were observed to be upregulated. A genomic region involved in adhesion was found to be highly induced in the biofilms of the S-layer containing strains *N. viennensis* and *N. maritimus*. This region included large adhesion proteins, approximately 1,000 to 3,000 amino acids in length, located next to other proteins potentially involved in adhesion, aggregation, and cell surface stability. Similar adhesion-related genes were not observed in *N. franklandianus*, which does not encode canonical S-layer genes^32^.

In both soil strains, the copCD/copC-copD gene region was highly upregulated with an additional putative cation channel protein in direct vicinity in *N. viennensis*. Specific to *N. franklandianus*, a highly expressed region of sialidases was observed.

TFBs were upregulated in all three species (Dataset S6). However, only in *N. maritimus* TFBs were highly upregulated and highly expressed (Fig. 6). Additionally, a secondary S-layer gene was identified to be upregulated in *N. maritimus* biofilms.

### Genetic distribution of bona fide biofilm features in AOA

To investigate the distribution of the identified bona fide biofilm genes amongst the diversity of AOA, a tree including 143 species was calculated and the presence of these genes in their genomes was analyzed (Fig. 7). MCOs exhibit considerable diversity, with clade specific patterns emerging. For instance, only the *Nitrososphaera* do not encode any MCO3s, while *Nitrosotalea* miss all MCOs except MCO3s. Genes encoding for adhesion proteins are completely absent from the *Nitrosocosmicus* clade, while they are commonly found in other soil lineages and in some marine lineages. Sialidases are highly enriched in all soil strains but are not commonly found in the marine strains. Most AOA encode a copC/copD variant, underlining their dependence on copper.

**Figure 7.**
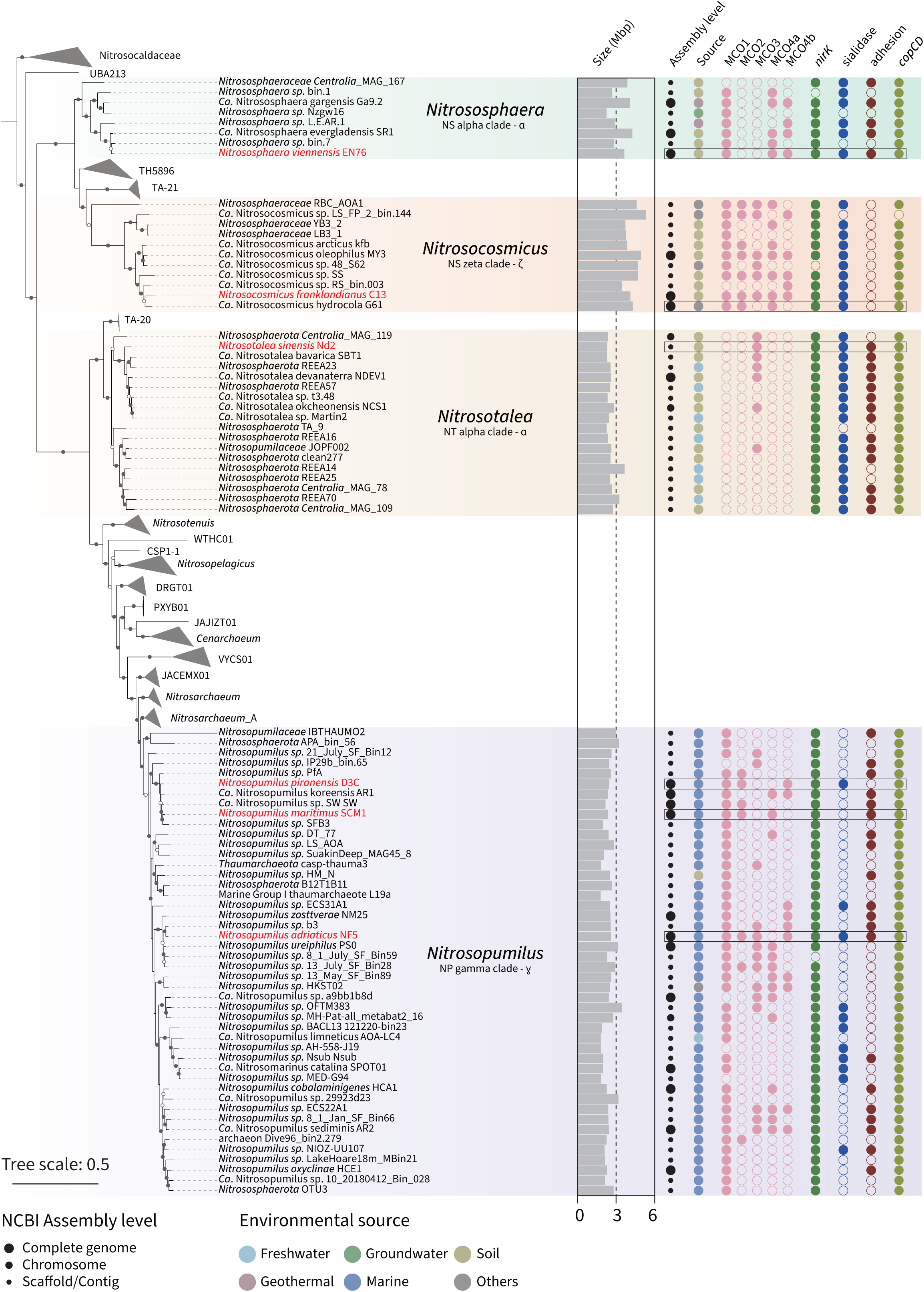
The distributions of genes of interest in genera *Nitrososphaera*, *Nitrosocosmicus*, *Nitrosotalea*, and *Nitrosopumilus*. The maximum likelihood phylogenomic tree of AOA was constructed based on a concatenation of 53 archaeal-specific markers (7589 columns, see Materials and Methods for details). Strains involved in this study are marked with red. Nodes with ultrafast bootstrap value ≥ 80% (60%) were indicated as solid (hollow) circles. The scale bar in the middle indicates 50% sequence divergence. The completeness and contamination of all shown genomes was ≥ 95% and < 5% respectively. Black circles represent different assembly levels according to NCBI. In brief, “complete genomes” are closed and contain no gaps, “chromosomes” are not closed and might still contain gaps while “scaffolds/contigs” are fragmented and are sure to contain gaps. An uncollapsed version of this tree including additional information is provided in the supplementary materials (Fig. S3).

## Discussion

In this study we systematically investigated the biofilm forming capabilities of AOA from diverse habitats and identified common as well as taxon-specific expression patterns under these growth conditions. All tested AOA formed biofilms without external triggers in optimal growth medium. Soil strains, especially *N. franklandianus,* exhibited higher capacities for biofilm formation, as expected from previous studies on members of the *Nitrosocosmicus* showing a higher genomic potential for biofilm formation^16,30,17^, or accelerated growth when surface associated^16^. Based on the presence of motility and chemotaxis genes, Bayer et. al. suggested that, although isolated from the same habitat, *N. piranensis* and *N. adriaticus* occupy different niches, floating in the water column or attached to marine snow/particles, respectively^33^. Our data support this hypothesis. The overall lower stability of biofilms from marine strains may be reflective of their environment, where communities attached to particles often undergo rapid successive changes^34^. Conversely, biofilms of terrestrial ecosystems might be more stable over prolonged periods of time.

### Attachment first

Two general strategies of biofilm formation were observed. *N. viennensis* and *N. maritimus* followed an attachment-first-strategy, gradually colonizing a surface, and subsequently building up a matrix. This strategy might be dependent on the presence of adhesion proteins, highly expressed in *N. viennensis* and *N. maritimus*. *N. piranensis* lacks these genes, potentially explaining its reduced biofilm-forming capabilities. Adhesion proteins are distributed irregularly among the different genomes of AOA hinting towards niche specialization by different AOA (Fig. 7). Several other genes are potentially involved in adhesion. For example, a DUF11 domain-containing protein (NVIE_000970) has been linked to cell surface stabilization during the aggregation of Methanothermobacter sp. CaT2^35^. Additionally, a putative Ca2+-binding protein (NVIE_001010) is present in the adhesion region. Calcium, known to enhance adhesion in *S. aureus* and *S. epidermidis*^36^, could similarly play a structural role in the biofilms of *N. viennensis*. In this context, a highly upregulated putative cation channel protein (NVIE_014290) may modulate Ca^2+^ levels of the biofilm. Adhesion proteins are recognized as key components of environmental biofilms^37^, and our findings suggest that these proteins are crucial for enabling the attachment strategy observed in these particular species.

### Aggregation first

Alternatively, *N. franklandianus* exhibited an aggregation-first-strategy. Cells deposited on the CG as suspended aggregates and three-dimensional structures were formed rapidly. Fittingly, it is common to observe cell aggregates in planktonic culture^19^, suggesting that *N. franklandianus* is growing as suspended aggregates. This appears to be a general feature of the *Nitrosocosmicus* clade as all enriched or isolated species were found to aggregate, or to be part of biofilms^19,16,38,39,17^. However, it is unclear how aggregates attach to surfaces, as specific mechanisms for attachment, such as adhesion proteins, are not present in the genomes (Fig. 7). Interestingly, canonical S-Layer genes are not found in the genus *Nitrosocosmicus*^40^. A genetic survey linked the absence of adhesion proteins to the genus *Nitrosocosmicus*, which lack an S-Layer (Fig. 7)^40^. Based on our observations of strong biofilm formation by *N. franklandianus* we propose biofilms as the preferred growth mode of this strain, and likely also of other members of the *Nitrosocosmicus* clade.

### The role of Multicopper Oxidases (MCOs) in biofilm formation

MCOs have been consistently observed genomically and transcriptionally in AOA^41,30,42^, although the biological function of most of them remains unknown. MCOs couple the oxidation of a substrate, that can be either organic or metal ions, to the reduction of oxygen to water^43^, but can have a wide range of substrates, sometimes even acting promiscuously^44^. MCOs have previously been speculated to be involved in the second step of ammonia oxidation as a functional homologue to bacterial HAO^41,45^, but the fact that MCOs are not conserved in all AOA genomes suggests that they are not involved in a central pathway^30^. The patchy distribution of different MCOs among AOA instead points towards diverse functions.

MCO1, 4a, and 4b were highly upregulated in *N. viennensis* under copper limitation, potentially oxidizing Cu+ to Cu2+ for bioavailability^42^, a known function for MCOs^46^. Contrastingly, in copper limited *N. maritimus,* the only upregulated MCO was of type MCO1^47^. Combining these observations, it is likely that MCO1 is indeed used for copper acquisition. Alternatively, MCO4a from *N. maritimus* has recently been shown to produce HNO from NH_2_OH which was proposed as a waste production pathway^48^. In this study, MCO4a was moderately to highly expressed in planktonic cultures and biofilms of all three species, and additionally highly upregulated in *N. viennensis* and *N. maritimus* biofilms (Fig. 6). MCO4a may therefore help cells to deal with nitrosative stress in biofilms where reactive molecules may be more concentrated. The upregulation of a nitroreductase in all strains supports an increase of nitro compounds that need to be detoxified^49^.

While MCO4a and MCO4b are overall similar, an alignment revealed that the T1 copper center in MCO4a is coordinated by histidines and a leucine, while the MCO4b copper center is coordinated by histidines and methionine (^42^, supplement). The methionine’s thioether group has been shown to modulate the redox potential of the coordinated CuII^50^ and might therefore be enabling a different function. The functional classification of MCOs using only primary structure is, however, significantly limited^44^. MCO4b and MCO3 were the most highly upregulated MCOs in biofilms of *N. viennensis* and *N. franklandianus* respectively. It is possible that these MCOs modify cell surfaces by oxidizing components like glycoproteins and in particular the glycosylated cell envelope. That MCO4b was the highest upregulated gene under both copper limitation and in biofilms of *N. viennensis* suggests a dual role of these cell surface modifications: (1) modifying cell surface structures to enable biofilm formation, and/or (2) aiding in the sequestration of positively charged ions, such as coppe*r*^51^. Additionally, MCOs in *N. franklandianus* were also highly expressed in planktonic conditions (Fig. 5.), which is in line with the observed continuous formation of cell aggregates in this strain. Several bona fide biofilm genes are also in close association with MCOs. The soil strains encode an “antibiotic biosynthesis monooxygenase” next to MCO4a and MCO3, respectively, indicating another oxygenase function that could modify the cell coat (rather than being involved in antibiotic biosynthesis). The presence of a putative electron carrier in all three strains, the ferredoxin/flavodoxin family protein, potentially indicates the importance of supplying electrons to MCOs (Fig. 6, Dataset S4).

### Transcription factor B as regulators in AOA biofilms

Transcription factors TBP and TFB, homologs of the eukaryotic basal transcription factors, are consistently found in all archaea and are essential for transcription. Similar to Halobacterium NRC-1, AOA encode several TFB proteins in their genomes that could be involved in global gene regulation^52^. Different from halobacteria, however, AOA possess only one TBP protein. While a switch in the most expressed TFB protein was observed in biofilm of *N. maritimus,* the regulation of soil strains seems to be more complex and involves the upregulation of one TFB in *N. viennensis* and three TFBs in *N. franklandianus* (Fig. 6, Dataset S6).

### Biofilm features specific to soil strains

The CopC/D transporters were upregulated by both soil strains. This may be necessary because the EPS matrix is known to sequester positively charged ions like copper^53^, trapping them and thus requiring increased activity of the uptake machinery. This was further supported by the presence of a cation channel protein located adjacent to copC/copD, which could supply other cations, such as Ca2+, that are bound by the biofilm matrix. It is plausible that copper acquisition genes in *N. maritimus* (CopD and MCO1) were not observed due to the single-layer architecture of the marine biofilm not impairing availability of copper.

Another shared feature of biofilms by soil strains is the upregulation of urease and urea transport genes in *N. viennensis* and *N. franklandianus* (Dataset S5) in the absence of urea. This might hint towards a broader substrate spectrum in biofilms. Cells in biofilms have been shown to employ highly efficient strategies for capturing diverse nutrients, surpassing that of free-living bacterial cells^26^.

A set of genes putatively involved in biofilm formation, including CAZymes, was previously proposed for *N. viennensis* and *N. franklandianus*^30,16^. Except for genes involved in adhesion, those genes were not differentially expressed in our datasets (Datasets S6). This might be in line with reports of polysaccharides sometimes being only a minor component in environmental biofilms, highlighting the importance of other factors like adhesion proteins^54^.

Sialidase encoding genes were upregulated in both soil strains, but prominently only in *N. franklandianus*. They have been shown to modify glycoproteins and glycolipids by cleaving sialic acid and are common compounds of environmental biofilms^55,56^. Furthermore, sialidase genes are present in the genomes of most AOA from soil and generally absent in the genomes of marine AOA (Fig. 7). Adjacent to the two sialidase genes in *N. franklandianus* is a gene for an electron carrier protein (ccd biogenesis protein, NFRAN_2155), which could be essential for supplying electrons to the sialidases, ensuring their activity and highlighting the importance of building up an EPS matrix.

### Specific responses in marine strain

Similar to *N. viennensis,* the marine strain *N. maritimus* relies on adhesion proteins for initial attachment. Instead of MCOs for surface modification, we observed a strong upregulation of an alternative S-layer protein that is likely important for biofilm formation and could possibly be needed to allow the introduction of the adhesion protein into the S-Layer.

Similarly, hemerythrin was found to be part of *N. maritimus* bona fide biofilm genes. Microbial communities associated with marine snow undergo complex successional changes^57^. While sinking, oxygen concentration decreases and hemerythrin could therefore act as an oxygen carrier^58^, similar to the situation in *M. capsulatus,* where bacteriohemerythrin protein has been shown to be highly upregulated under low oxygen conditions^59^.

## Conclusion

In conclusion, we have demonstrated that all tested AOA have the capacity to form biofilms, although to different extents. They rely heavily on cell surface modifications and/or adhesion, for which two strategies were observed (Fig. 8). Distinct expression patterns illustrate that mostly individual solutions have evolved for the shared growth mode of biofilm formation in AOA, probably driven by the different ecological niches.

**Figure 8.**
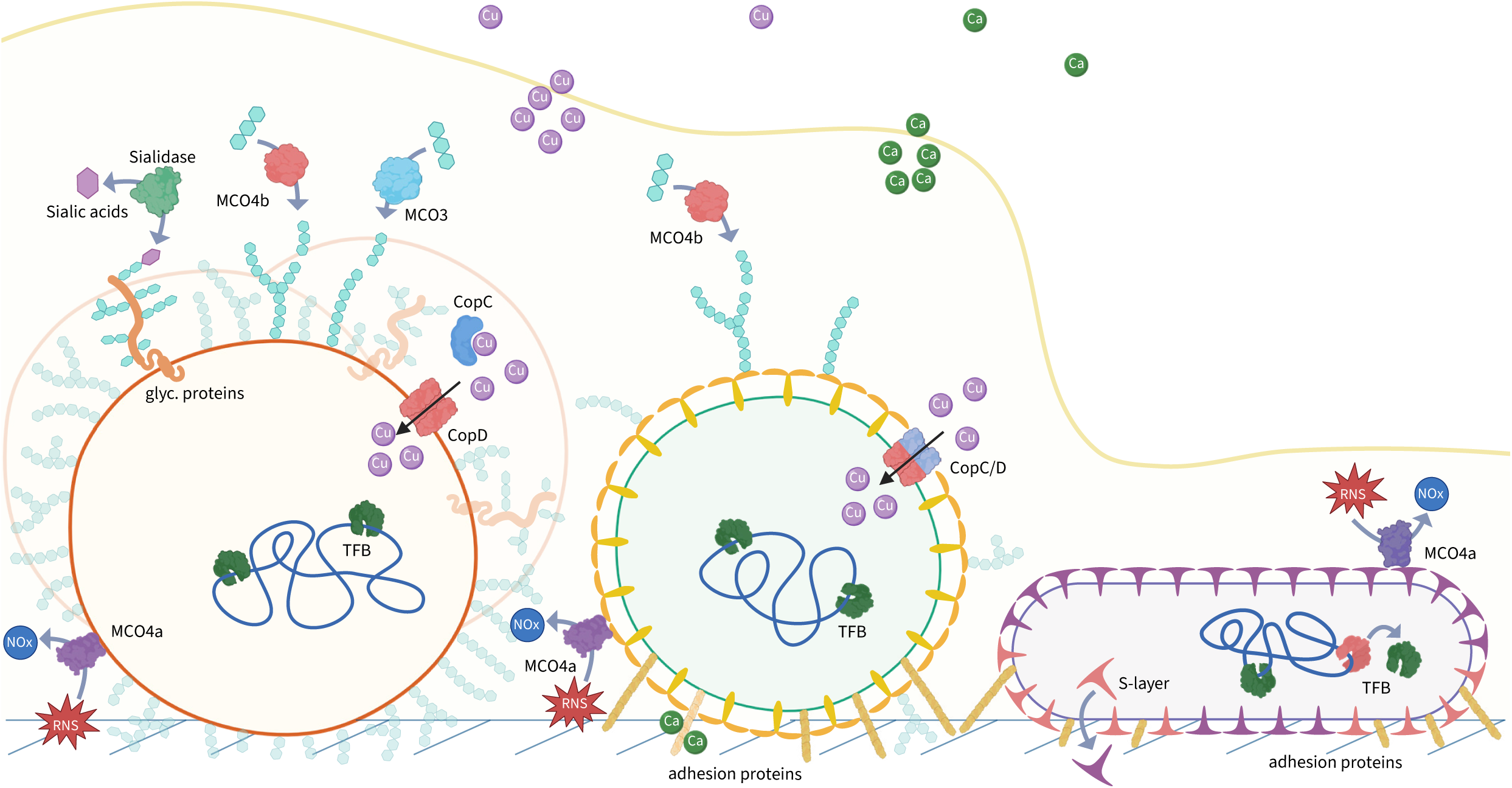
Schematic overview of AOA biofilms highlighting a subset of functional genes and their putative functions. Represented genes were chosen based on the common themes emerging from the bona fide biofilm gene analysis shown in Fig. 6: Adhesion, Cell surface modification including, Strong upregulation and expression of MCOs, Copper acquisition machinery and transcriptional regulation.

AOA are among the few archaea globally distributed with great ecological success. This is likely in part due to their high energy efficiency, encoding the most efficient CO_2_ fixation pathway^60^ and their high affinities for ammonia^10,61^. Their broad ecological success could, however, also be determined by their ability to form biofilms. From this work, it is clear that to understand the role of AOA in the environment, they must also be studied in the context of biofilms. This is especially true for AOA derived from soils and may even be imperative for the genus *Nitrosocosmicus*. This study sets the stage for future investigations of the biofilm structures and of the activities and ecophysiological effects of AOA in biofilms.

## Acknowledgments

We thank Andrea Malits and Sarah Harrer for their excellent technical support, Daniela Gruber for her expertise and help with scanning electron microscopy and Dr. Melina Kerou for discussions. Microscopy was performed at the Core Facility Cell Imaging and Ultrastructure Research, University of Vienna - member of the Vienna Life-Science Instruments (VLSI). The RNA sequencing was performed by the Next Generation Sequencing Facility at Vienna BioCenter Core Facilities (VBCF), member of the Vienna BioCenter (VBC), Austria. The computational results of this work have been achieved using the Life Science Compute Cluster (LiSC) of the University of Vienna. This project was funded by the Austrian Science Fund, Projects P36287 (The Ammonia Oxidation Process in Archaea) and Z437 (Archaea Ecology and Evolution), as well as EU Horizon 2020 twinning project ActionR (Research Action Network for Reducing Reactive Nitrogen Losses from Agricultural Ecosystems) No. 101079299.

