## Supplementary_material for "Biofilm lifestyle as a common trait of ammonia-oxidizing archaea"

### **Material and Methods**

All datasets generated during the current study are available in the pribasnig repository, under following link: <https://www.ncbi.nlm.nih.gov/sra/PRJNA1156597>

The whole-transcriptome data generated in this study have been deposited in the NCBI BioProject database under the accession project number (PRJNA1156597), Submission ID: SUB14703162.

#### *Strains*

*Nitrososphaera viennensis* EN76 was isolated from garden soil^1^. *Ca.* Nitrosocosmicus franklandianus C13 was isolated from arable soil and kindly provided by Dr. Graeme Nicol (Ecole Centrale de Lyon)^2^. Nitrosotalea sinensis Nd2 was isolated from acidic paddy field soil and kindly provided by Dr. Graeme Nicol (Ecole Centrale de Lyon)^3^. *Nitrosopumilus maritimus* SCM1 was isolated from the sediment of a tropical marine aquarium and kindly provided by Dr. Martin Könneke (University of Washington, Seattle)^4^. *Nitrosopumilus piranensis* D3C and *Nitrosopumilus* *adriaticus* NF5 were both isolated from coastal marine surface waters and kindly provided by Dr. Barbara Bayer (University of Vienna)^5^.

#### *Culture maintenance*

Cultures of the terrestrial AOA *N. viennensis*, *N. franklandianus* and *N. sinensis* were grown in freshwater medium (FWM) buffered with HEPES or MES, while the marine AOA *N. maritimus*, *N. adriaticus and* *N. piranensis* were grown in synthetic crenarchaeota medium (SCM) as described previously. For a detailed description of FWM and SCM refer to Reyes et al., 2020, Table 1 and Könneke et al., 2005 respectively^4,6^. All cultures were supplemented with ammonia and grown in 30 ml polystyrene containers (Greiner Bio-One, G201170) holding 20 ml medium in the dark. AOA were incubated at and supplemented with following temperatures and ammonia concentrations: *N. maritimus*, *N. adriaticus and N. piranensis* [1 mM ammonia, 28°C]. *N. sinensis* [1 mM ammonia, 35°C]. *N. viennensis*, *N. franklandianus* [2 mM ammonia, 42°C]. The pH of FWM buffered with HEPES or MES was between 7.5-7.7 and 5.0-5.3 respectively. The pH of unbuffered SCM media was between 7.3-7.5. Media were prewarmed before inoculation. *N. viennensis* and *N. franklandianus* were shaken at 80 r.p.m. to improve oxygen diffusion. *N. viennensis* was grown without the addition of vitamins and with 1 mM pyruvate to scavenge reactive oxygen species hydrogen peroxide (H_2_O_2_) otherwise inhibiting growth^7^. For detailed descriptions of each species full medium and standard growth conditions refer to the above cited isolation papers and supplementary table S1. The growth of all stock cultures was followed by measuring nitrite concentrations colorimetrically via the Griess reaction as described previously^6^. Final concentrations of all media constituents are summarized in Table S1. Stock cultures were routinely checked for contamination microscopically and by spotting culture aliquots on R2A agar plates.

#### *RNA extraction*

Total RNA was extracted from five biological replicates of frozen microscopy slides (MS) containing *N. viennensis*, *N. franklandianus* and *N. maritimus* biofilms and five planktonic batch controls, grown in 125mL species-specific medium in 250mL Schott bottle flasks (Schott). Cells were harvested via filtration (MCE Membrane Filter, 0.2 µm pore size, 47mm, Merck) during mid exponential growth. Following the filtration step, filters were inserted into 15 mL falcon tubes (Fisher Scientific), immediately frozen on dry ice and stored at -70°C. RNA of *N. viennensis* and *N. maritimus* samples was extracted from filters and MS using the mirVana™ miRNA Isolation Kit (Thermo Fisher) according to the manufacturer’s instructions. The lysis procedure was adjusted for *N. franklandianus* as follows. After resuspending cells in the Lysis/binding buffer, *N. franklandianus* cells were transferred to a screw tube containing lysis matrix B (0.1 mm silica spheres, MP Biomedicals) and mechanically homogenized three times using a FastPrep-24™ Classic Instrument (MP Biomedicals) at 4.0 m/s, 25s, cooling samples on ice for 5 minutes after each run. The RNA extraction was continued as described in the manual, finally eluting RNA in 50-100µl of DEPC. Concentrations and possible contaminations were measured by Nanodrop (NanoPhotometer N60, Implen). If necessary, phenol was removed in an additional cleanup step as follows. The volume of the samples was adjusted to 300µl with DEPC. After adding 30µl 3M Na-Acetate and 1µl Glycogen, samples were vortexed, 550µl 100% EtOh was added and samples incubated on ice for 1 hour. RNA was pelleted by centrifugation at 11kxg, 20min, 4°C, liquid discarded and RNA washed with 70% EtOH. After removing the liquid and air-drying the pellet for 10 min, RNA was resuspended in 50-100µl of DEPC. Finally RNA was DNAse digested (with TURBO™ DNase (Thermo Fisher) for 1h. To exclude any traces of DNA were still left in the samples, a PCR was done using mco4_b (NVIE_019250) primer^8^, or amoA primer^9^ as previously described.

#### *Transcriptomic analysis*

RNA was submitted to the Vienna Biocenter Facility (VBC) for Illumina NovaSeq S4 PE150 XP sequencing with a species specific rRNA depletion step. Raw reads were downloaded from VBC, checked using md5sum and are available online under the link provided above. FastQC v.0.12.1^10^ and multiQC v.1.25^11^ were used to initially analyze the data. Reads were trimmed using fastp v.0.23.4^12^ with following specific settings: --trim_front1 12 --trim_poly_g --detect_adapter_for_pe --average_qual 30 --length_required 30. rRNA reads were sorted out with SortmeRNA^13^, and remaining mRNA reads were mapped to the specific genomes using HISAT2 v.2.2.1^14^ and counted using featureCounts^15^. Raw counts are available for all datasets. All further analyses were conducted in R-Studio version 2021.09.1+372^16^, and samples from biofilms were compared against planktonic samples as controls. All scripts are available on github under: https://github.com/pribasnig/Biofilm_NV_NF_NM. Counts were analyzed for differential expression using DESeq2 employing the default settings^17^. The Benjamini Hochberg DESeq2 default step was used to control for false discovery rate (FDR). Genes with an adjusted p-value <0.01 and a log2-FC in biofilms of ≥ 1.0 or ≤ -1.0 were considered significantly up/downregulated. For principal component analysis, data was normalized using the rlog function in DESeq2. A PCA plot was then produced from the normalized data using the R packages ggplot2 (Wickham, 2016) and ggrepel^18^. Conditions were tested for significant differences using PERMANOVA analysis from the R package vegan^19^. TPM values were calculated by dividing the read counts by the length of each gene in kilobases, yielding reads per kilobase (RPK). All RPK values per sample were summed and divided by 1,000,000 giving a per-million scaling factor. Finally RPK values were divided by the scaling factor to yield transcripts per million (TPM). Log2 of TPM was calculated to use for color coding of TPM values. Conservation of genes between *N. viennensis*, *N. franklandianus* and *N. maritimus* was analyzed by attributing protein families to AOA genes as previously described by^20^. A list of AOA genomes used can be found in Dataset S1. Once assigned, conservation of protein families between the species was compared using the package dplyr in R^21^. Based on their protein families, genes significantly upregulated in all possible combinations of the 3 species were identified (NV+NF+NM, NV+NF, NV+NM, NF+NM, NV, NF, NM). A venn diagram was produced with the R package VennDiagram^22^, displaying the results. To further investigate the genes important under growth in biofilms, the top 50 genes in log2FC were further filtered to also be amongst the top 100 TPM BF. Genes that failed the defined cutoffs, but that were in the same genomic region and followed the same general trends of upregulation and expression as above defined genes were manually added for further analysis.

#### *Phylogenetic analysis*

The candidate sequences of MCO were identified via hmmsearch v3.4^23^ (PF07281 and PF07282) against the AOA representative genomic dataset (completeness ≥ 95% and contamination < 5%), and then aligned with MAFFT v7.526^24^. Poorly aligned regions with ≥ 95% gaps were filtered with TrimalAL v1.5^25^ and the maximum-likelihood phylogeny for the MCO alignment was inferred using the IQ-TREE v2.3.4^26^ with 1000 UFBoot replicates. The atypical sequences were removed manually. All trees were uploaded to iTOL for visualization.

The 53 archaeal-specific concatenated ribosomal proteins identified with GDTB-Tk v2.4.0^27^ were chosen to generate phylogenetic trees of AOA. Except for family UBA213 (*Nitrososphaerota* SAT137 with completeness 84.3%), only the representative genomes with completeness ≥95% and contamination ≤5% were retained as the final dataset. Poorly aligned regions were removed using TrimAL v1.4 with the parameters “-gt 0.95 –cons 50”. The maximum-likelihood phylogeny for the concatenated alignment was inferred using the IQ-TREE v2.3.4 with 1000 UFBoot replicates. The best-fitting protein model, Q.insect+F+R8, was determined using ModelFinder.

### **Supplementary tables and figures**

| **Media composition of *N. viennensis*, *N. franklandianus*, *N. sinensis*, *N. maritimus*, *N. adriaticus* and *N. piranensis*** | | | | |
| --- | --- | --- | --- | --- |
|  | Final concentration [mM] | | | |
|  | *N. viennensis* | *N. franklandianus* | *N. sinensis* | *N. maritimus, N. adriaticus & N. piranensis* |
| **Salts [autoclaved]** |  |  |  |  |
| NaCl | 17 | 17 | 17 | 444.9 |
| MgCl_2_ 6H_2_O | 1.9 | 1.9 | 1.9 | 24.59 |
| CaCl_2_ 2H_2_O | 6.8 | 6.8 | 6.8 | 10.2 |
| KCl | 6.7 | 6.7 | 6.7 | - |
| NaHCO_3_ | 2 | 2 | 2 | 1 |
| MgSO_4_ 7H_2_O | - | - | - | 20.28 |
| KBr | - | - | - | 0.84 |
| KH_2_PO_4_ | 1.14 | 1.14 | 1.14 | * |
| **Nutrients [filtered]** |  |  |  |  |
| NH_4_Cl | 2 | 2 | 1 | 1 |
| **Buffer (pH) [filtered]** |  |  |  |  |
| HEPES (7.5) | 10 | 10 | - | - |
| MES (5) | - | - | 10 | - |
| **Trace Elements [filtered]** |  |  |  |  |
| FeNaEDTA solution | 0.75 | 0.75 | 0.75 | 0.75 |
| Non-chelated trace elements |  |  |  |  |
| HCl | 0.1 | 0.1 | 12.5 | 0.1 |
| H_3_BO_3_ | 5*10^-4^ | 5*10^-4^ | 5*10^-4^ | 5*10^-4^ |
| MnCl_2_ 4H_2_O | 5*10^-4^ | 5*10^-4^ | 5*10^-4^ | 5*10^-4^ |
| CoCl_2_ 6H_2_O | 8*10^-4^ | 8*10^-4^ | 8*10^-4^ | 8*10^-4^ |
| NiCl_2_ 6H_2_O | 1*10^-4^ | 1*10^-4^ | 1*10^-4^ | 1*10^-4^ |
| ZnSO_4_ 7H_2_O | 5*10^-4^ | 5*10^-4^ | 5*10^-4^ | 5*10^-4^ |
| Na_2_Mo_7_O_24_ 4H_2_O | 1.5*10^-4^ | 1.5*10^-4^ | - | 1.5*10^-4^ |
| (NH_4_)_6_ Mo_7_O_24_ 4H_2_O | - | - | 2.9*10^-5^ | - |
| CuCl_2_ H_2_O | 1.4*10^-5^ | 1.4*10^-5^ | 1.4*10^-5^ | 1.4*10^-5^ |
| **Vitamins [filtered]** |  |  |  |  |
| Vitamin solution | - | + | - | - |
| Biotin |  | 8*10^-5^ |  |  |
| Folic acid |  | 5*10^-5^ |  |  |
| Pyrodoxine HCl |  | 4.8*10^-4^ |  |  |
| Thiamine HCl |  | 1.4*10^-4^ |  |  |
| Riboflavin |  | 1.3*10^-4^ |  |  |
| Nicotinic acid |  | 4*10^-4^ |  |  |
| DL Panthothenic acid |  | 2*10^-4^ |  |  |
| p-aminobenzoic acid |  | 3.6*10^-4^ |  |  |
| Choline chloride |  | 1.4*10^-3^ |  |  |
| Vitamin B12 |  | 7.3*10^-9^ |  |  |
| Antibiotics [filtered] |  |  |  |  |
| Kanamycin (µg/ml) | 8.5*10^-2^ (50) | 8.5*10^-2^ (50) | - | - |
| Other [filtered] |  |  |  |  |
| Na-pyruvate | 1 | - | - | - |
| KH2PO4* | - | - | - | 2.29 |
| Growth conditions & transfer criteria | | | | |
| Temperature [°C] | 42 | 42 | 35 | 28 |
| Shaking [rpm] | 80 | 80 | - | - |
| Stock inoculation [%v/v] | 0.25 | 1 | 2 | 5 |
| Nitrite at transfer [mM] | 1.2-1.8 | 1.2-1.8 | 0.15-0.25 | 0.5-0.8 |

* KH_2_PO_4_ was sterile filtered and added separately for marine AOA strains

Table S1. **Growth conditions of AOA species used in this study.**

| **Biomass accumulation rate** | | | | | |
| --- | --- | --- | --- | --- | --- |
| **Transfer** | ***N. viennensis*** | ***N. franklandianus*** | ***N. maritimus*** | ***N. adriaticus*** | ***N. piranensis*** |
| 1 | 0.61 | 0.93 | 0.80 | 0.83 | 0.46 |
| 2 | 0.79 | 1.09 | 1.02 | 1.07 | 0.43 |
| 3 | 0.90 | 1.18 | 0.98 | 1.22 | 0.44 |
| 4 | 0.95 | 1.73 | 1.30 | 1.35 | 0.53 |
| 5 | 1.27 | 2.92 | 1.68 | 1.79 | 0.54 |
| 6 | 1.42 | 3.44 | 1.38 | 2.01 | 0.64 |
| 7 | 1.68 | 4.14 | 1.25 | 2.15 | 0.72 |
| 8 | 1.89 | 3.83 | 1.22 | 1.73 |  |
| 9 | 2.32 | 4.33 | 1.22 | 1.53 |  |
| 10 | 2.20 |  | 1.28 | 1.40 |  |
| 11 | 2.53 |  |  | 1.15 |  |
| 12 |  |  |  | 1.03 |  |

Table S2: **Biomass accumulation rates per transfer per organism.**

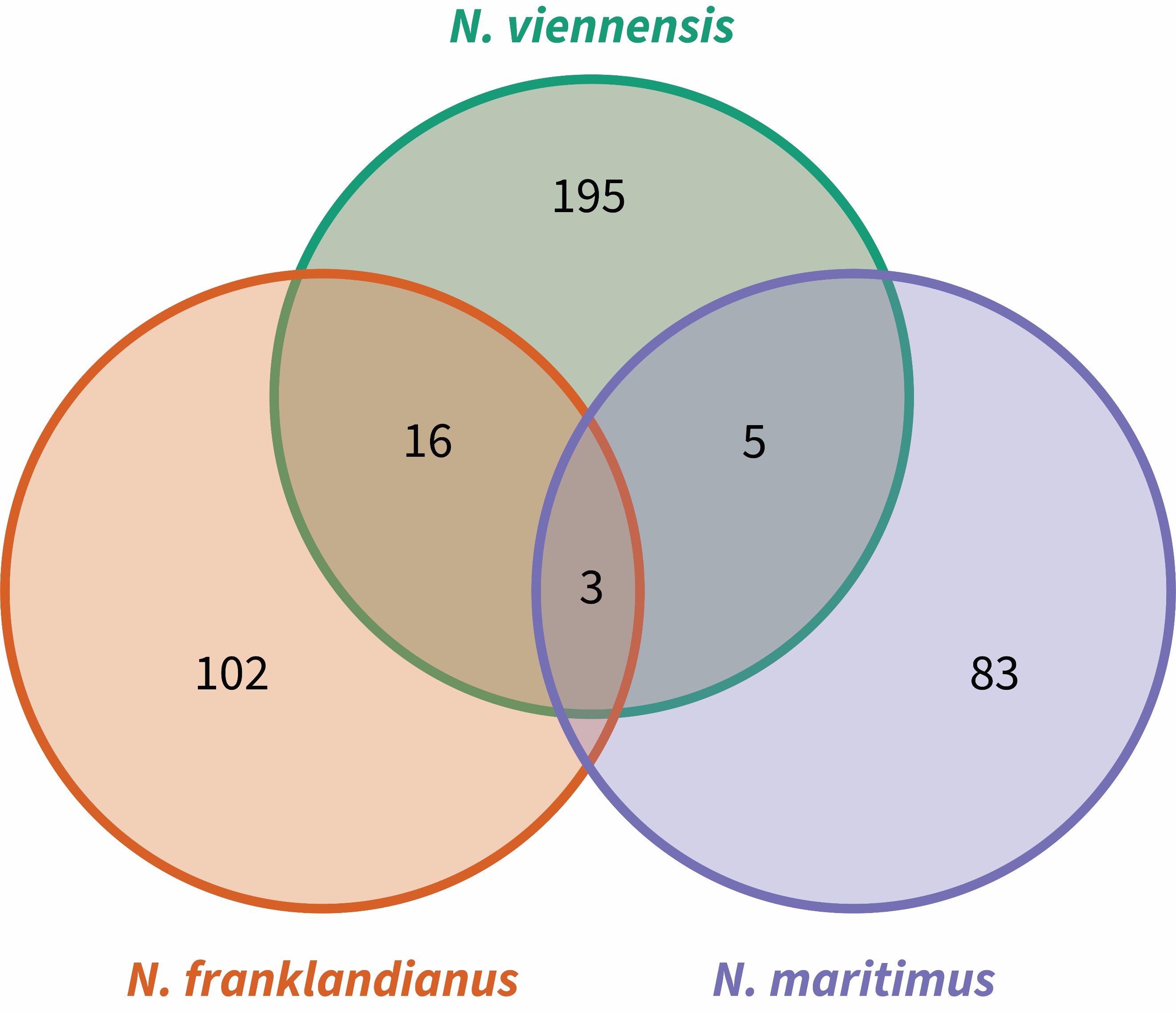

Supplementary Figure S1: **Venn Diagram of shared protein families of significantly upregulated genes in biofilms (log2FC >1, padj <0.001).** The Venn Diagram shows shared or unique protein families between N. viennensis, N. franklandianus and N. maritimus in all possible combinations.

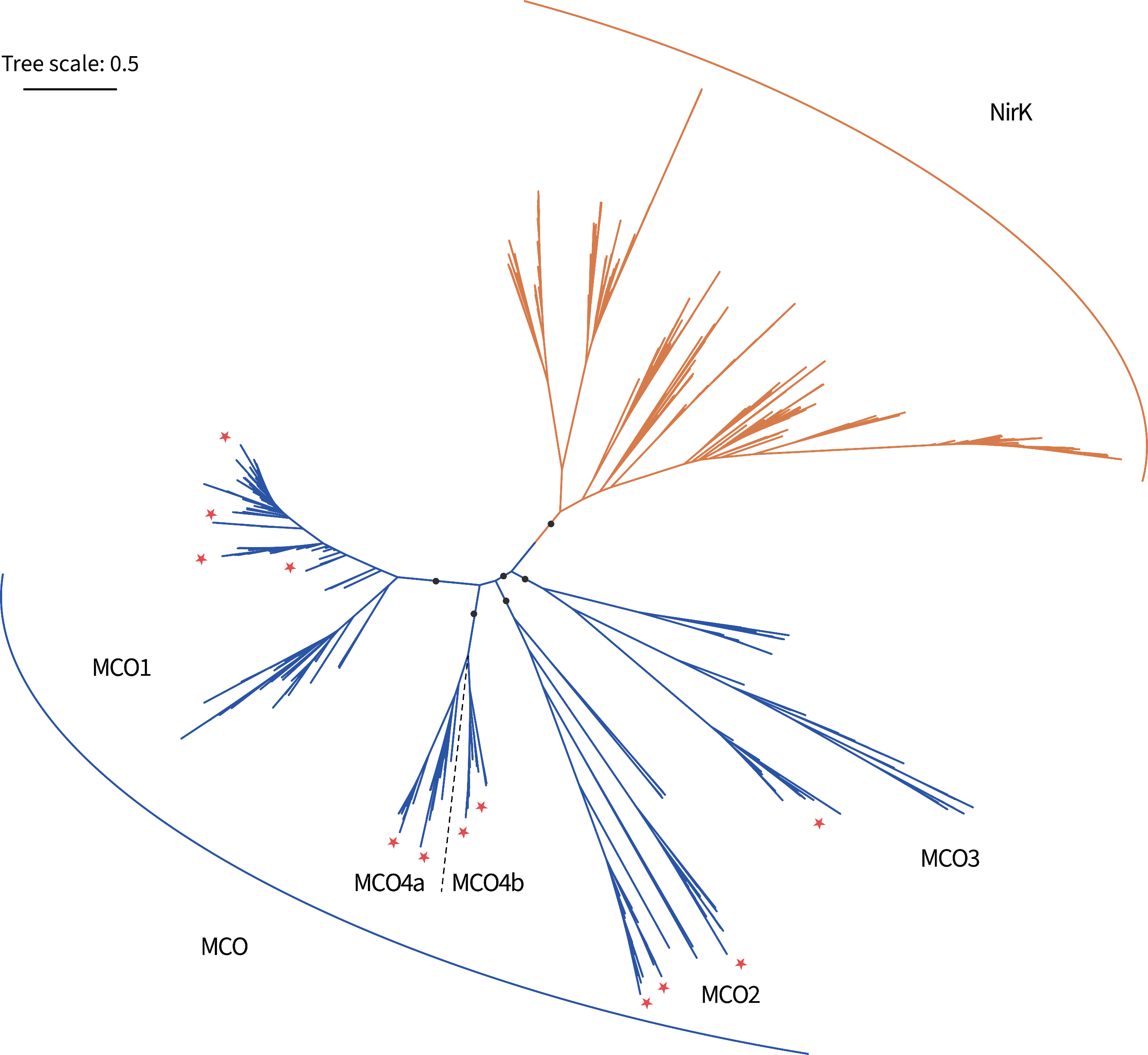
Supplementary Figure S2: **The full unrooted tree for NirK and multicopper oxidases (MCO1, MCO2, MCO3, MCO4a, and MCO4b).** NirK and MCO sequences are shown in orange and blue respectively. The placements of MCOs of organisms included in this study are marked with red asterisks. Nodes with ultrafast boot strap ≥ 80% are indicated as solid circles and the scale bar indicates 50% sequence divergence.

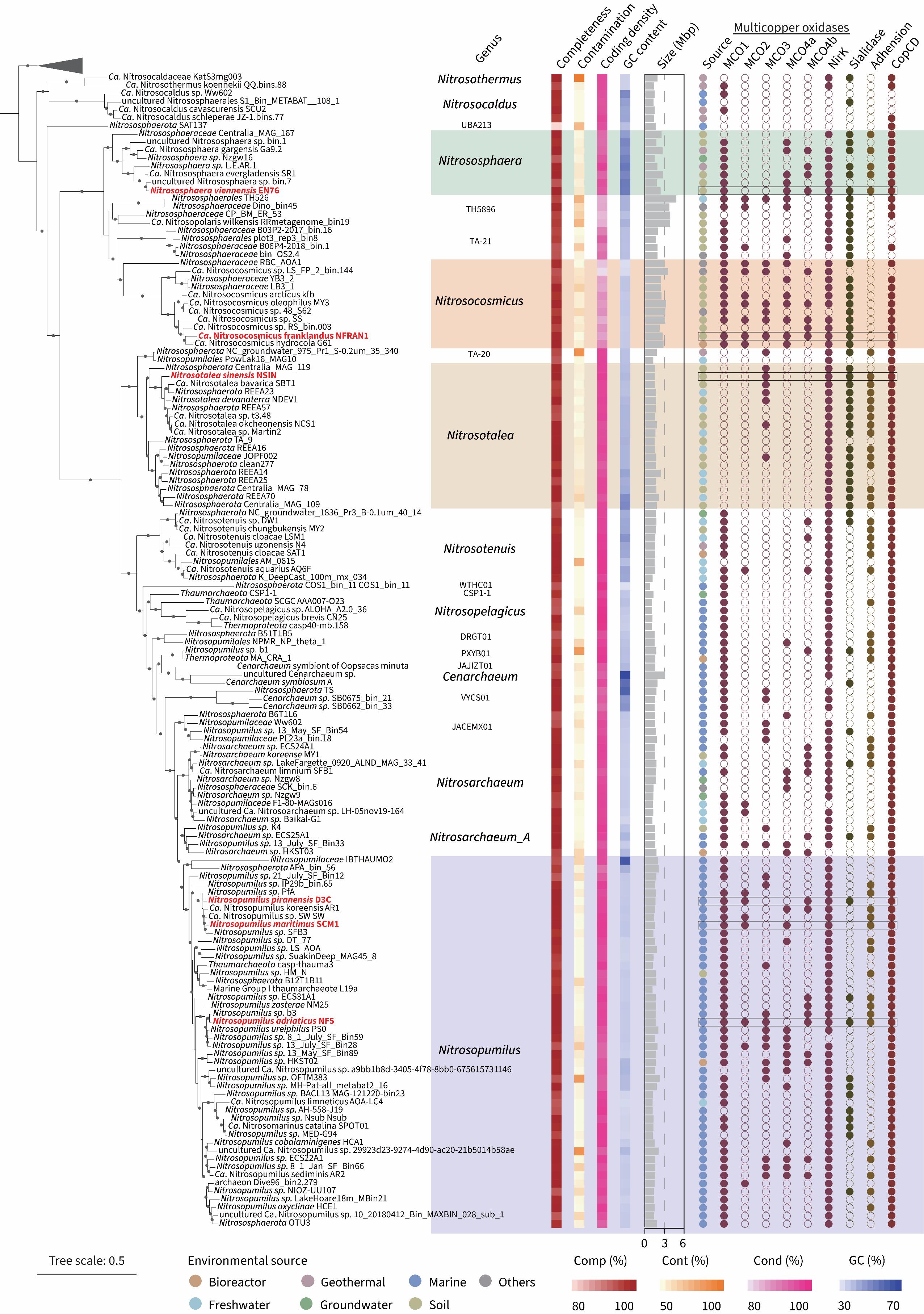

Supplementary Figure S3: **The overall distribution of genes of interest in AOA.**

**Supplementary Data**

**Dataset_S1_growth_raw_data_and_curves.xlsx**

This excel file contains raw growth data in the form of nitrite measurements of all 6 strains used in this study, used for figure 1.

**Dataset_S2_Nviennensis.xlsx**

This excel file contains all transcriptomic data for *N. viennensis*, raw counts, DeSeq output and calculated TPMs for all genes. Additionally, significantly differently expressed genes, and the Top25 differently expressed genes used in figure 4 are displayed here.

**Dataset_S3_Nfranklandianus.xlsx**

This excel file contains all transcriptomic data for *N. franklandianus*, raw counts, DeSeq output and calculated TPMs for all genes. Additionally, significantly differently expressed genes, and the Top25 differently expressed genes used in figure 4 are displayed here.

**Dataset_S4_Nmaritimus.xlsx**

This excel file contains all transcriptomic data for *N. maritimus*, raw counts, DeSeq output and calculated TPMs for all genes. Additionally, significantly differently expressed genes, and the Top25 differently expressed genes used in figure 4 are displayed here.

**Dataset_S5_conserved_protein_families_Venn_diagramm.xlsx**

This excel file contains the conserved protein families between *N. viennensis, N. franklandianus* and *N. maritimus* in all possible combinations, used in supplementary figure S1.

**Dataset_S6_figures_discussion_data.xlsx**

This excel file contains transcriptomic data from *N. viennensis, N. franklandianus* and *N. maritimus* used in figures 5&6, all conserved upregulated protein families, CAZymes, and species used for calculating protein families.
