## Supplementary figures and images for "Biofilm lifestyle as a common trait of ammonia-oxidizing archaea"

### Supllementary_figure_S2

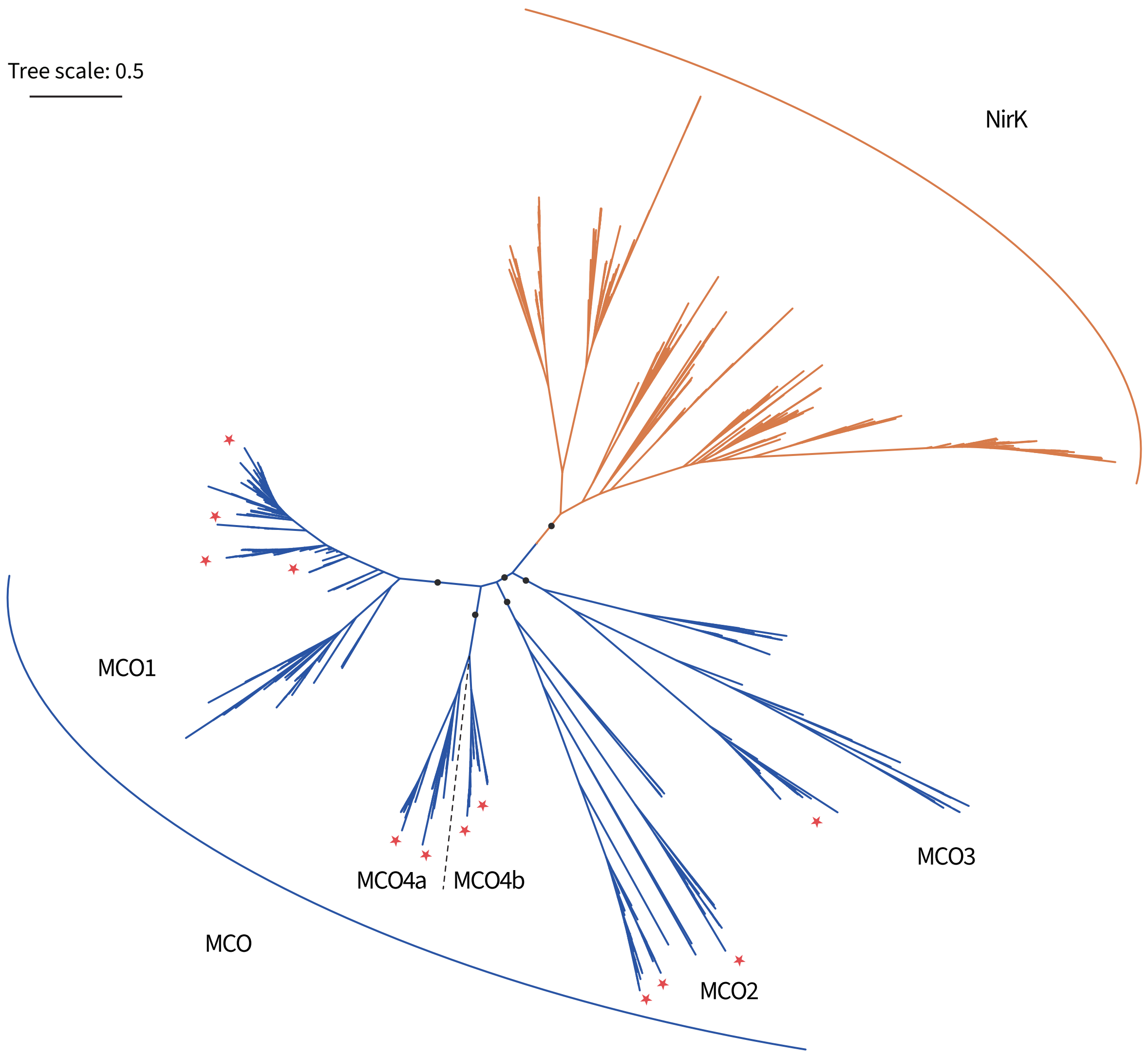

### Supplementary_figure_S1

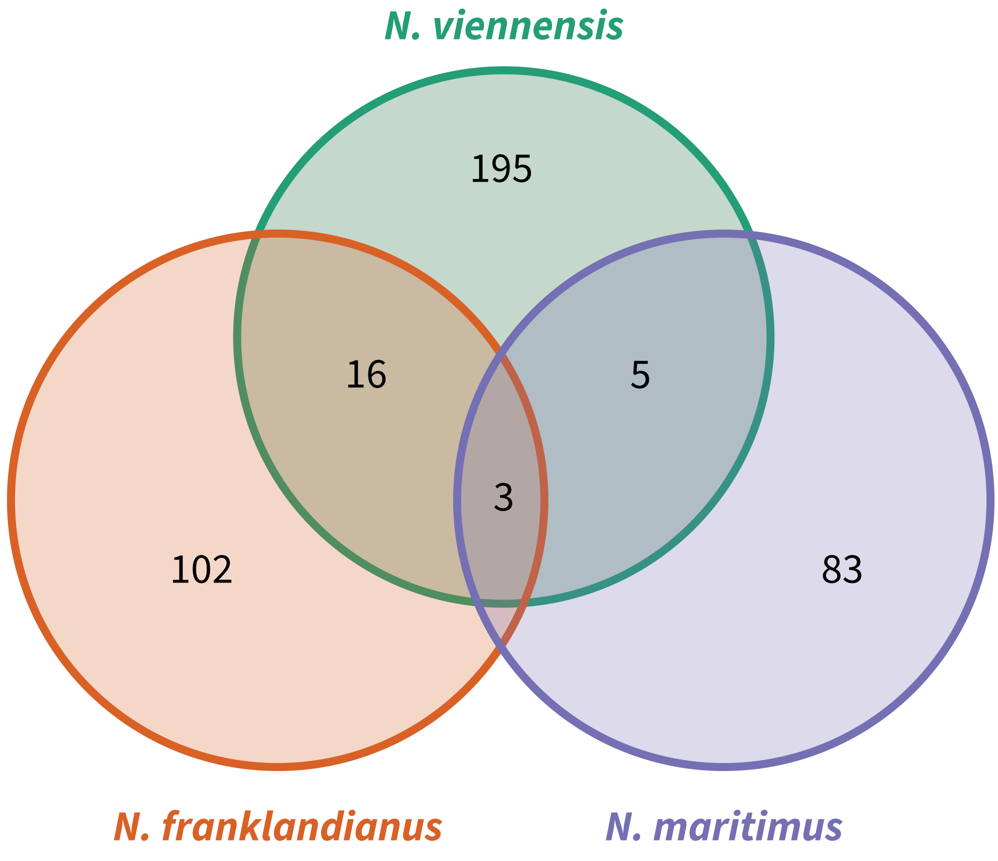

### Supplementary_figure_S3

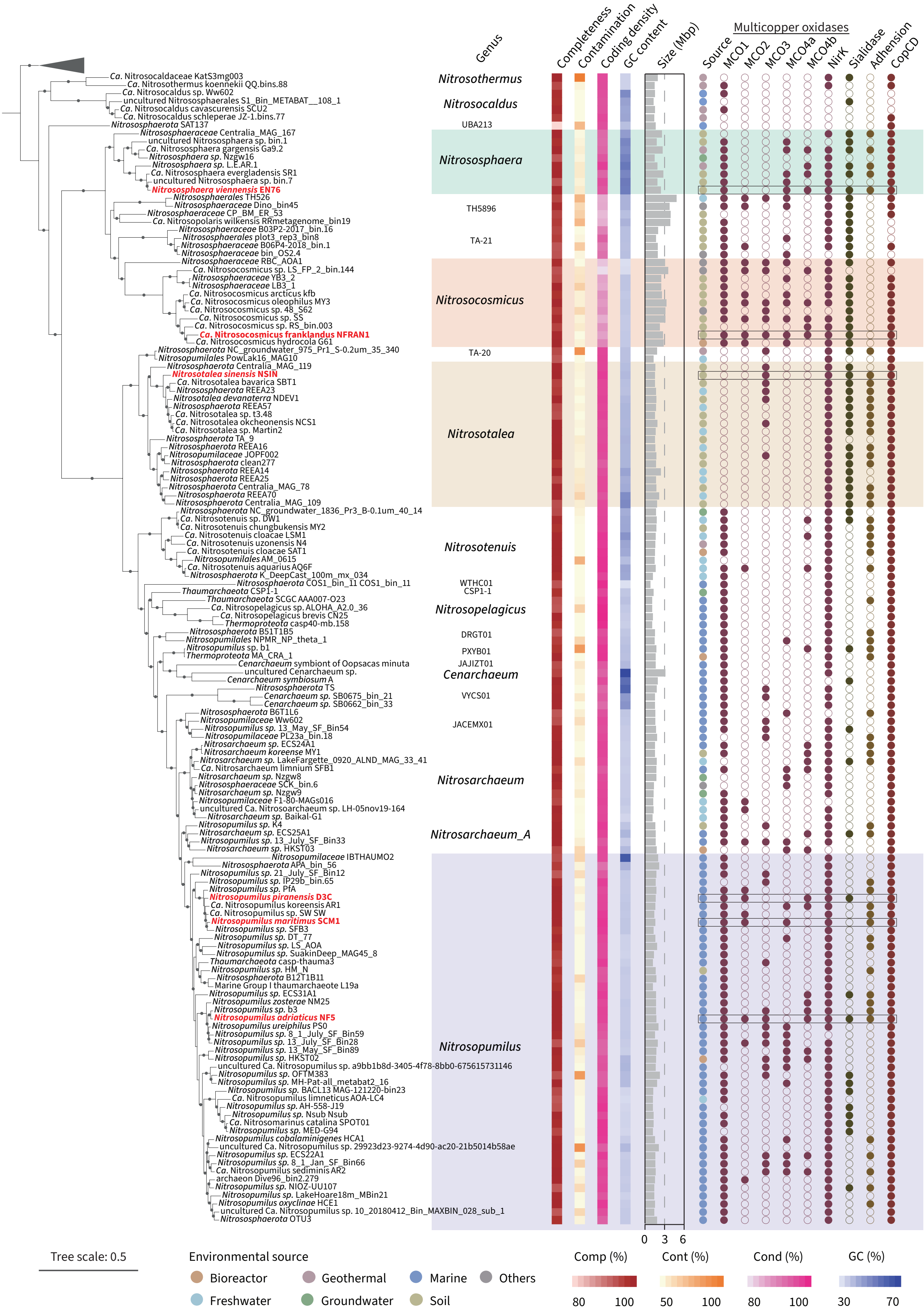
